# Impact of intensive care unit admission during handover on mortality: a propensity matched cohort study

**DOI:** 10.1101/813279

**Authors:** Thais Dias Midega, Newton Carlos Viana Leite Filho, Antônio Paulo Nassar, Roger Monteiro Alencar, Antônio Capone Neto, Leonardo Rolim Ferraz, Thiago Domingos Corrêa

## Abstract

**Introduction:** Handover is a process of transferring information, responsibility and authority for providing care of critically ill patients from a departing intensivist to an oncoming intensivist. The effect of i admission during a medical handover on clinical outcomes is unknown.

**Objectives:** Our purpose was to evaluate the impact of ICU admission during a medical handover on clinical outcomes.

**Methods:** Post hoc analysis of a cohort study addressing the effect of ICU admissions during the handover on outcomes. This retrospective, single center, propensity matched cohort study was conducted in a 41-bed open general ICU located in a private tertiary care hospital in São Paulo, Brazil. Based on time of ICU admission, patients were categorized into two cohorts: handover group (ICU admission between 6:30 am to 7:30 or 6:30 pm to 7:30 pm) or control group (admission between 7:31 am to 6:29 pm or 7:31 pm to 6:29 am). Patients in the handover group were propensity matched to patients in the control group at 1:2 ratio. Our primary outcome was hospital mortality.

**Results:** Between June 1, 2013 and May 31, 2015, 6,650 adult patients were admitted to the ICU. Following exclusion of ineligible participants, 5,779 patients [389 (6.7%) in handover group and 5390 (93.3%) in control group] were eligible for propensity score matching, of whom 1,166 were successfully matched [389 (33.4%) handover group and 777 (66.6%) in control group]. Before matching, hospital mortality was 14.1% (55/389 patients) in handover group compared to 11.7% (628/5,390) in control group (p=0.142). After propensity-score matching, ICU admission during handover was not associated with increased risk of ICU (OR, 1.40; 95% CI, 0.92 to 2.11; p=0.11) and hospital (OR, 1.23; 95%CI, 0.85 to 1.75; p=0.26) mortality. ICU and hospital length of stay did not differ between the groups.

**Conclusion:** In this propensity-matched single center cohort study, ICU admission during medical handover did not affect clinical outcomes.

## Introduction

The demand for intensive care unit (ICU) beds is increasing worldwide, reflecting an increased life expectancy and prevalence of chronic conditions [1]. Since the availability of ICU beds is limited, it is crucial to improve ICU organizational and operational characteristics to improve efficiency and outcomes. In Brazil, most of the ICUs have a full time, i.e., 24/7, in house ICU physician (intensivists) coverage [2]. As a result, frequently transitions of care between health care professionals are expected to happen.

Handover is a process of transferring information, therapeutic plan and responsibility of ICU patients from a departing to an oncoming provider [3]. Handover process is specially challenging at the ICU, owing to the complexity of critically ill patients and how fast their clinical conditions may change [4]. Additionally, patients admitted to the ICU are usually unstable, demanding timely resuscitative maneuvers, invasive procedures and therapeutic interventions during the first hours of ICU admission [5]. Therefore, one could expect that ICU admission during the handover process, when crucial information to the care of critically ill patients are being transferred form one intensivist to the other, might be associated with an increased incidence of communication failures, medical errors, unexpected adverse events and worsened clinical outcomes [6].

Several studies have shown an association between deficient handovers and clinical outcomes [7-9]. For instance, a study conducted at the emergency department (ED) found that inadequate handover adversely affected approximately 5% of patients, resulting in delayed therapy [10]. A review of malpractice claims in the ED evidenced that 24% of missing diagnosis were caused by inadequate handovers [11]. In another study, patients submitted to cardiac surgery who received intraoperative handover of anesthesia care had a 43% higher chance of hospital mortality compared to patients that did not received intraoperative handover [12]. The leading cause of ineffective handover is related to communications problems, such as omissions or corrupted information, mostly due to distractions. Therefore, it is of paramount importance conduct the handover processes in a quiet and free of interruptions environment [4].

## Objective

Our primary objective was to evaluate the effect of ICU admission of critically ill patients during a medical handover on hospital mortality in a tertiary care hospital. Secondary objectives were to compare resource use and clinical outcomes among patients admitted and patients not admitted to the ICU during the handover period.

## Methods

### Study design and settings

The present study is a post hoc analysis of a retrospective single center cohort study that investigated the impact of ICU readmission on resource use and clinical outcomes [13]. The original study and this post hoc analysis were approved by the Local Ethics Committee at Hospital Israelita Albert Einstein with waiver of informed consent (CAAE:54065716.3.0000.0071).

### Setting

This study was conducted in a private tertiary care hospital in São Paulo, Brazil, comprising 662 inpatient beds, one adult general, open model ICU with 41 beds and 91 step-down unit beds.

### Patients

All consecutive patients aged ≥18 years old admitted to the ICU between June 1, 2013 and May 31, 2015 were included. Patients with missing core data [age, gender, time of ICU admission, ICU admission diagnosis, Simplified Acute Physiology score (SAPS 3 score) at ICU admission, ICU and hospital length of stay (LOS) and vital status at hospital discharge were excluded (Fig 1).

**Fig. 1.**
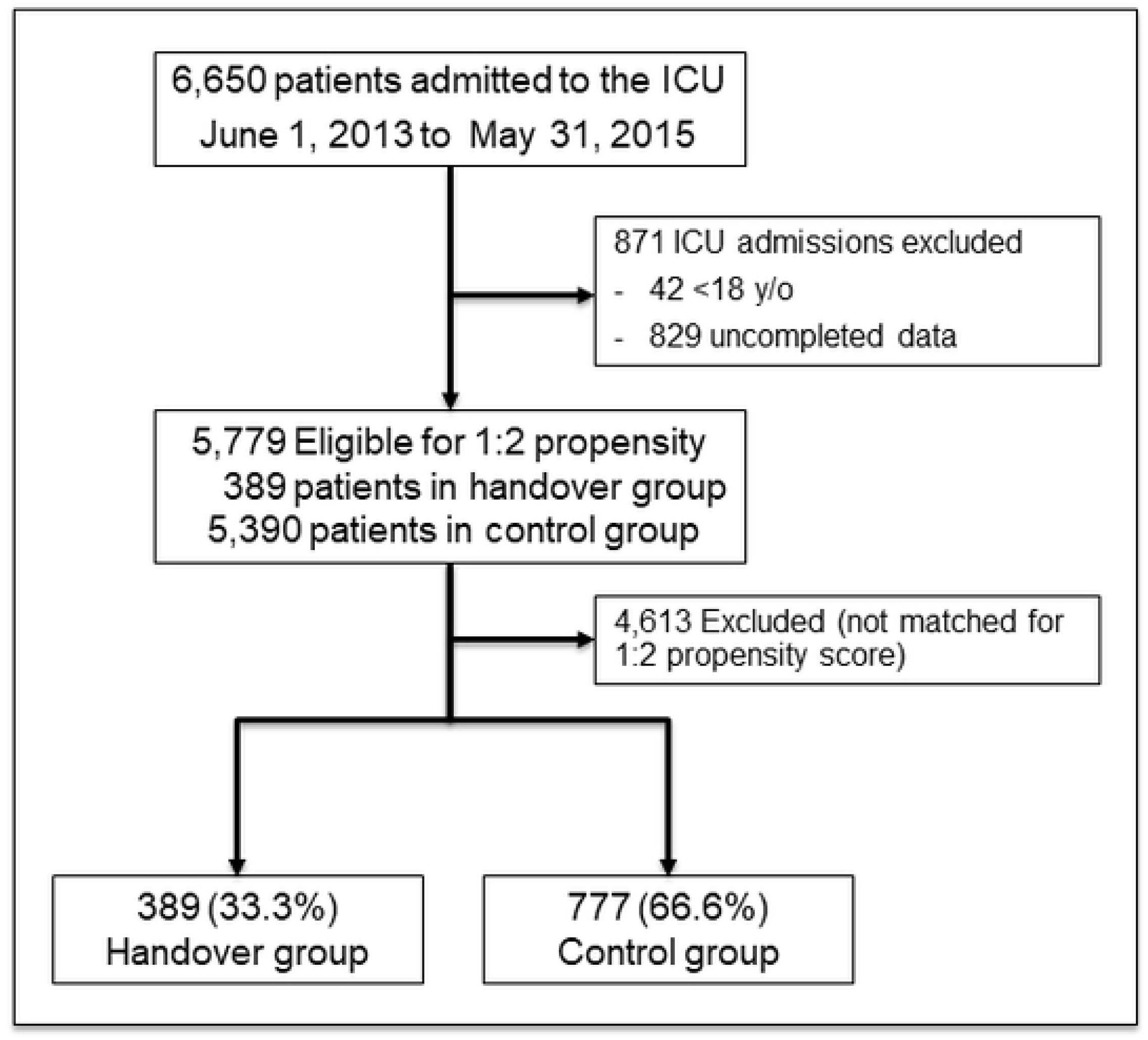
Patient flow charge.

### Data collection and study variables

All study data were retrieved from Epimed Monitor System^®^ (Epimed Solutions, Rio de Janeiro, Brazil), which is an electronic structured case report form, where patients data are prospectively entered by trained ICU case managers [14]. Collected variables included demographics, comorbidities, location before ICU admission, time of ICU admission and discharge, reason for ICU admission, SAPS 3 score at ICU admission [15], ICU admission diagnosis, need for invasive support [need for vasopressors, mechanical ventilation, noninvasive mechanical ventilation (NIV) and renal replacement therapy (RRT)] on ICU admission and during ICU stay, destination at ICU discharge, ICU and hospital LOS, frequency of ICU readmission, hospital mortality, mortality at ICU discharge and at day 90.

### Definitions

Handover was defined as the transfer of care between a departing intensivist to an oncoming intensivist. The handover time was defined as the time period between 6:30 am to 7:30 am and between 6:30 pm to 7:30 pm. Based on time of ICU admission, patients were categorized into two cohorts: handover group (ICU admission between 6:30 am to 7:30 or 6:30 pm to 7:30 pm) or control group (admission between 7:31 am to 6:29 pm or 7:31 pm to 6:29 am).

### ICU characteristics and the handover process

On-duty ICU physicians are available 24 hours a day at a rate of one intensivist per every ten beds. There is no reduction in personnel or in ICU activities during night shifts or at weekends. Multidisciplinary clinical rounds involving ICU physicians, nurses, respiratory therapists, nutritionists, psychologist and clinical pharmacists are performed daily. ICU admissions are made by on-duty intensivists, whereas discharge is a consensus decision-making process involving on-duty intensivists and the primary physician, i.e., the physician who will accept the patient outside the ICU [13].

The handover process was performed twice daily, at 7:00 am and 7:00 pm, when the departing intensivist shift ends. The departing ICU physicians usually begin to prepare for the handover process 30 minutes prior to the handover time. The handover process may last up to 60 minutes. Orally and written (standardized handover tool) medical-handover were performed for every ICU patient.

Handover between oncoming and departing ICU nurses / respiratory therapists occurred at the same time as the handover between intensivists.

### Statistical analysis

Categorical variables are presented as absolute and relative frequencies. Continuous variables are presented as median with interquartile ranges (IQR). Normality was assessed by the Kolmogorov-Smirnov test. Comparisons were made between the handover group and control group. Categorical variables were compared with chi-square test or Fisher exact test when appropriate. Continuous variables were compared using independent t test or Mann-Whitney U test in case of non-normal distribution.

We used a propensity-score matching designed to mitigate confounding by accounting for differences in the patient’s characteristics. Propensity scores for handover group were estimated for each patient with logistic regression using seventeen relevant patients characteristics (age, gender, SAPS III score, reason for index ICU admission, index admission source, presence of systemic hypertension, diabetes mellitus, cancer, congestive heart failure, chronic obstructive pulmonary disease, chronic kidney disease, chronic kidney disease requiring long-term dialysis and liver cirrhosis, use of vasopressor, renal replacement therapy and mechanical ventilation during index ICU stay). Patients with missing data were excluded from the database. Based on the propensity score weighted estimators we constructed a propensity score-matched cohort. Matching was performed using nearest neighbor matching without replacement, with each patient in handover group matched to two patients in the control group. A caliper width of 0.10 of the standard deviation of the logit of the propensity score was used for the development of matching [16, 17].

Statistical tests were 2-sided, and a p<0.05 was considered statistically significant. No adjustment for multiplicity was applied across the analyses. Statistical analyses were performed using IBM^®^ SPSS^®^ Statistics version 22.0 for Windows. Plots were performed using GraphPad Prism version 6.00 for Windows (GraphPad Software, California, USA).

## Results

### Characteristics of studied population

Between June 2013 and July 2015, 6,650 patients were admitted to ICU. After exclusion of 871 ICU admissions due to incomplete core data and/or age under eighteen, 5,779 patients were included in the analysis [389 (6.7%) in handover group and 5,390 (93.3%) in control group]. The median (IQR) age of the cohort was 67 (53-80) years and 56.5% of patients were men. From the 5,779 eligible patients for propensity score matching, 1,166 were successfully matched. Out of those, 389 (33.4%) were in handover group and 777 (66.6%) were in control group (Fig.1). A histogram showing the distribution of hours of ICU admission for the study population (n=5,779 patients) is shown in fig. 2.

**Fig 2.**
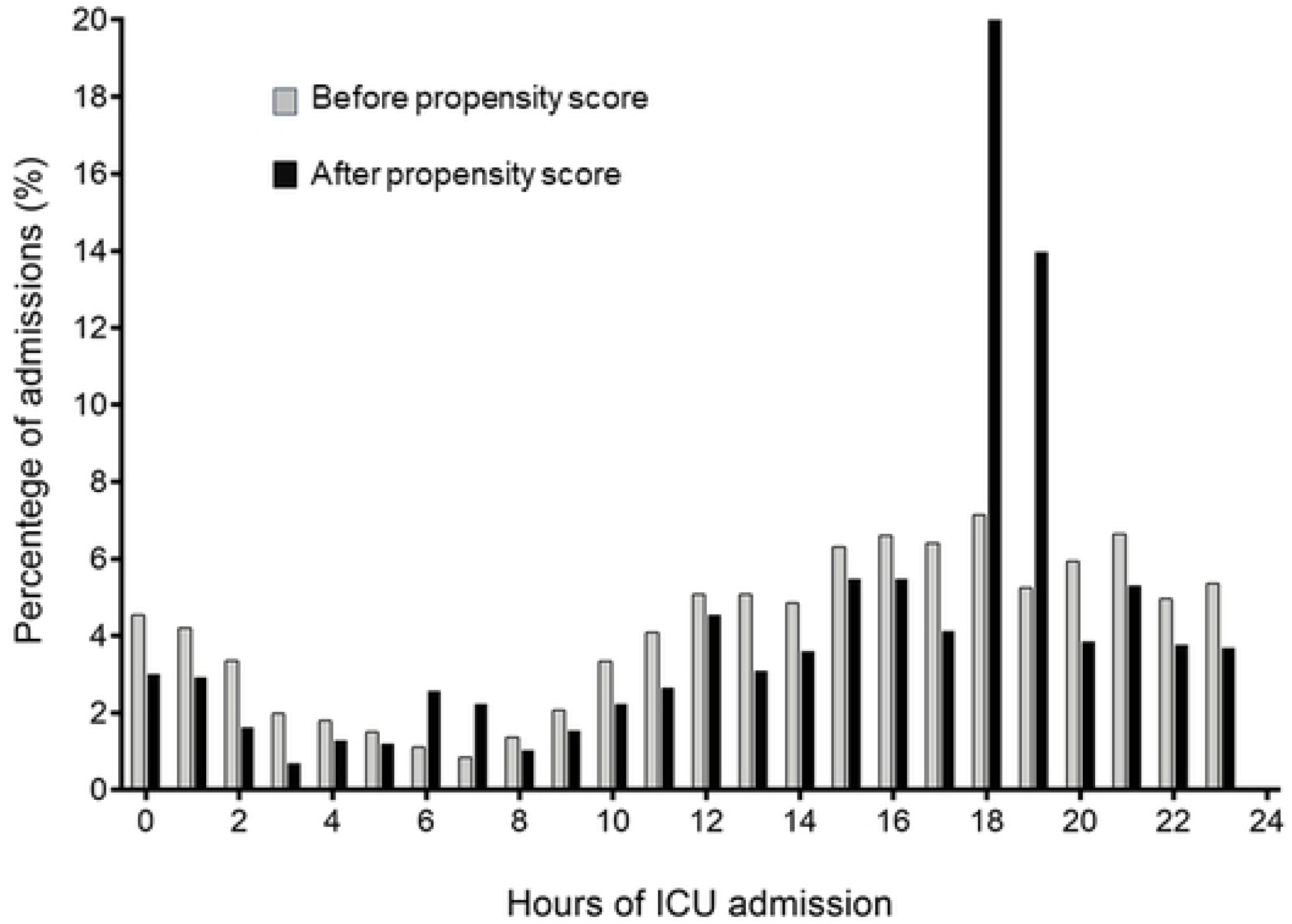
Percentage of admissions to the ICU per hours of the day.

### Cohort before propensity score matching

Before propensity score matching, age, gender, SAPS III reason for ICU admission, admission source, admission diagnosis, frequency of co-morbidities, as well as support need on ICU admission, such as mechanical ventilation, non-invasive ventilation, RRT and vasopressors, were similar between patients in the handover group and control group (S1 Table).

Hospital mortality was 14.1% (55/389 patients) in handover group compared to 11.7% (628/5,390 patients) in control group (OR, 1.25; 95%CI, 0.92 to 1.68; p=0.142). Resource use, expressed as the need of vasopressors, mechanical ventilation, non-invasive ventilation and RRT did not differ between handover group and control group. Length of ICU and hospital stay and frequency of ICU readmissions were also similar among the groups (Table 1).

**Table 1.**
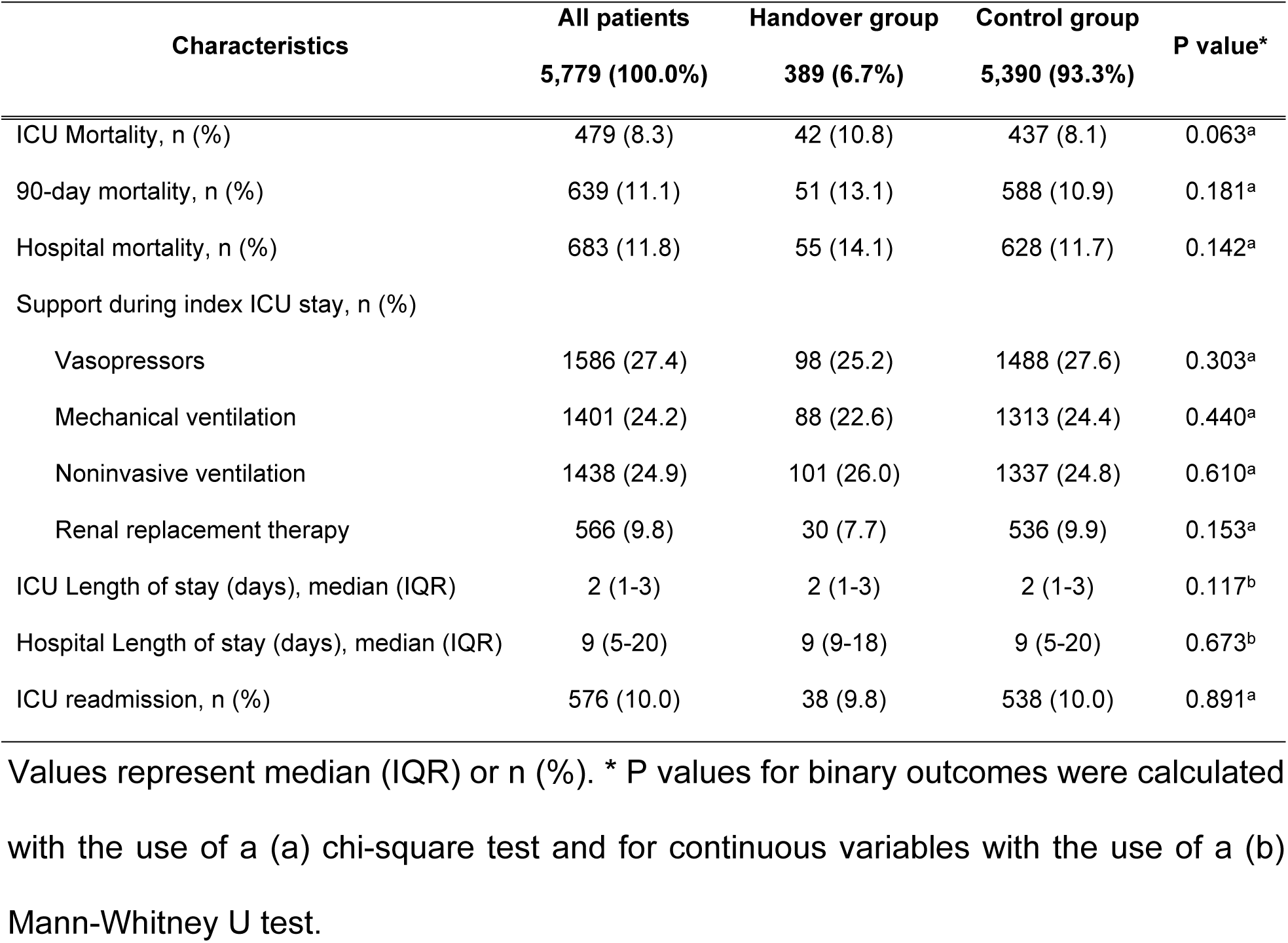
Outcomes before propensity score matching.

### Cohort after propensity score matching

The propensity-matched cohort had a median (IQR) age of 67 (53-80) years, 54.9% (640/1,166) of patients were men with a median (IQR) SAPS III score of 43 (32-55) (Table 2). The study groups were well balanced with respect to age, gender, SAPS III score at index ICU admission, reason for index ICU admission, admission source, prevalence of co-morbidities, admission diagnosis and the need of supportive therapy on index ICU admission (Table 2).

**Table 2.**
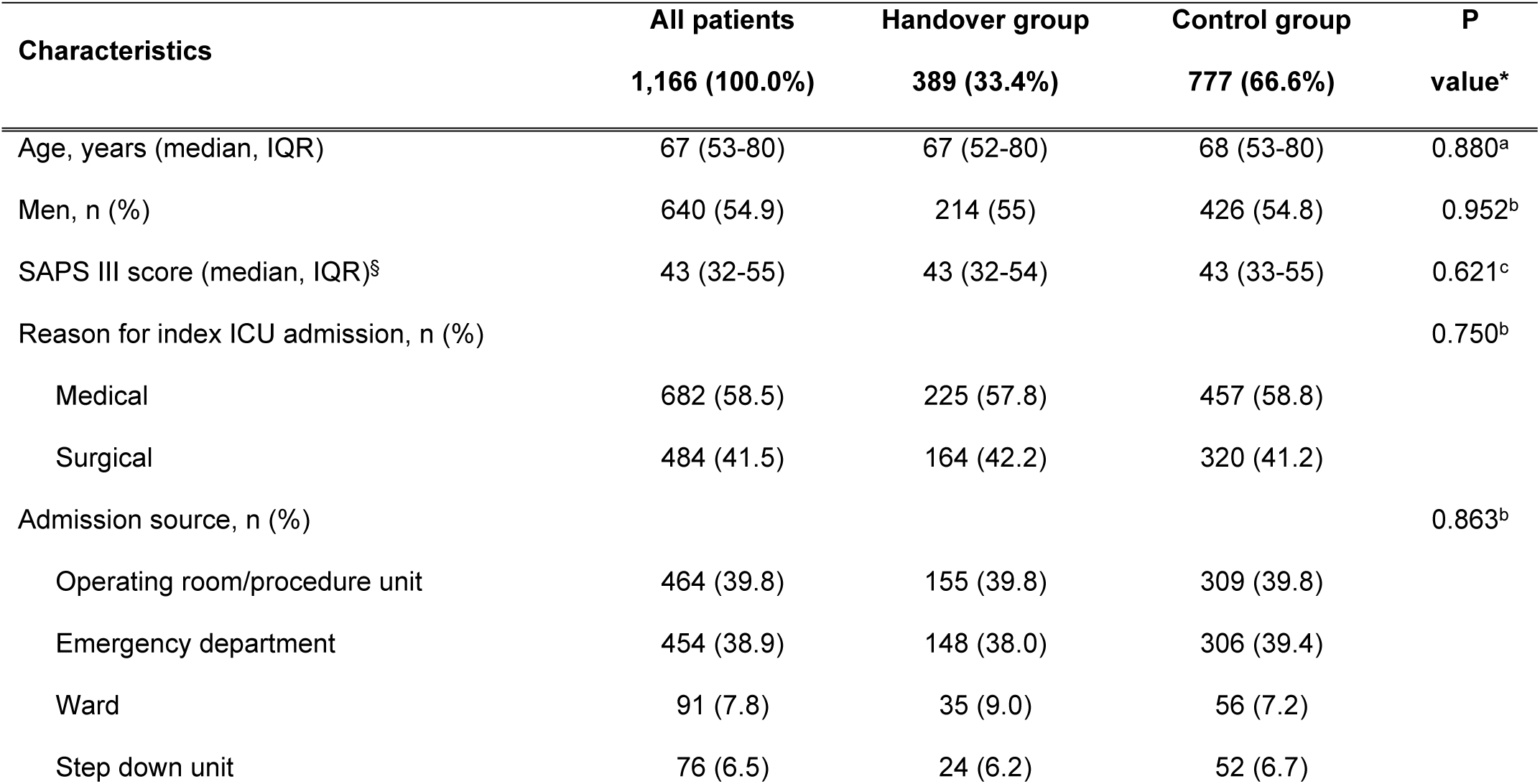
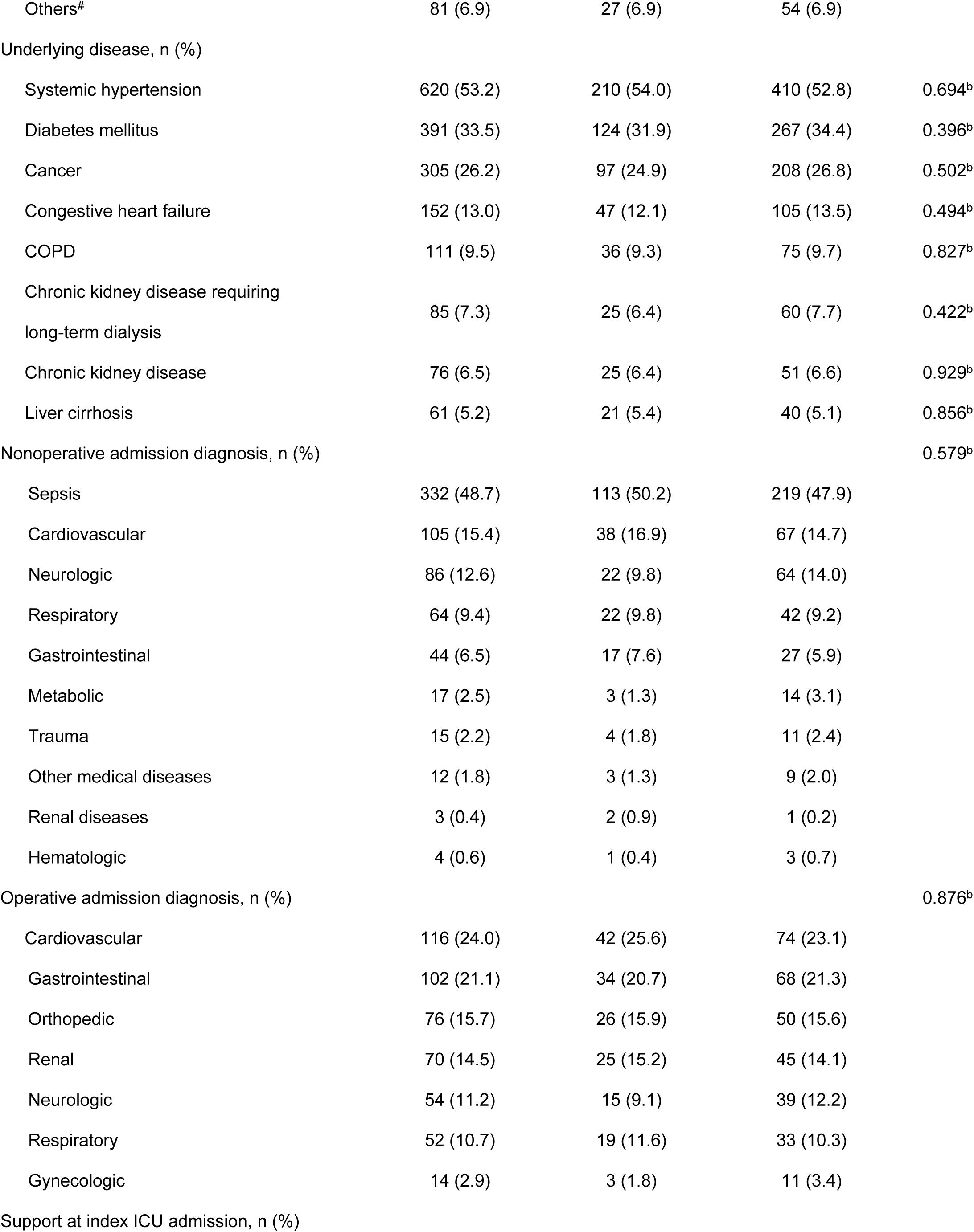
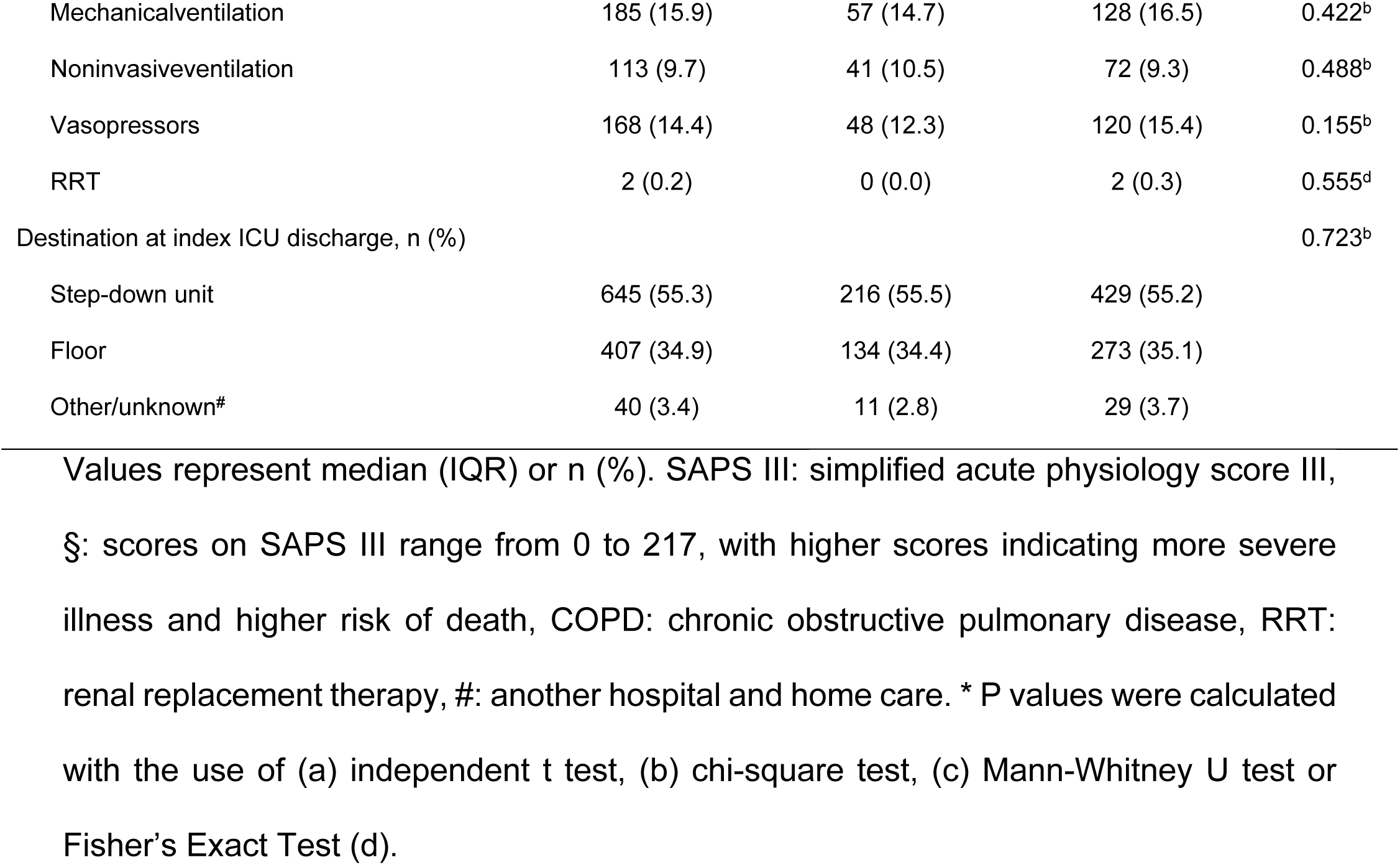
Baseline characteristics of study participants after propensity score matching.

Hospital mortality was 14.1% (55/389 patients) in handover group compared to 11.8% (92/777 patients) in the control group patients (OR, 1.23; 95%CI, 0.85 to 1.75; p=0.265). The use of vasopressors, mechanical ventilation, non-invasive ventilation and renal replacement therapy, ICU and hospital length of stay and the frequency of ICU readmissions did not differ between groups (Table 3).

**Table 3.** Outcomes after propensity score matching.

| Characteristics | All patients<br>1166 (100.0%) | Handover group<br>389 (33.4%) | Control group<br>777 (66.6%) | P value* |
| --- | --- | --- | --- | --- |
| ICU Mortality at ICU, n (%) | 104 (8.9) | 42 (10.8) | 62 (8.0) | 0.111<sup>a</sup> |
| 90-day mortality, n (%) | 139 (11.9) | 51 (13.1) | 88 (11.3) | 0.375<sup>a</sup> |
| Hospital mortality, n (%) | 147 (12.6) | 55 (14.1) | 92 (11.8) | 0.265<sup>a</sup> |
| Support during index ICU stay, n (%) |  |  |  |  |
| Vasopressors | 302 (25.9) | 98 (25.2) | 204 (26.3) | 0.696<sup>a</sup> |
| Mechanical ventilation | 270 (23.2) | 88 (22.6) | 182 (23.4) | 0.760<sup>a</sup> |
| Noninvasive ventilation | 321 (27.5) | 101 (26.0) | 220 (28.3) | 0.397<sup>a</sup> |
| Renal replacement therapy | 94 (8.1) | 30 (7.7) | 64 (8.2) | 0.756<sup>a</sup> |
| ICU Length of stay (days), median (IQR) | 2 (1-4) | 2 (1-3) | 2 (1-4) | 0.258<sup>b</sup> |
| Hospital Length of stay (days), median (IQR) | 10 (5-20) | 9 (5-18) | 10 (5-20) | 0.689<sup>b</sup> |
| ICU readmission, n (%) | 120 (10.3) | 38 (9.8) | 82 (10.6) | 0.678<sup>a</sup> |

Values represent median (IQR) or n (%). \* P values for binary outcomes were calculated with the use of a (a) chi-square test and for continuous variables with the use of a (b) Mann-Whitney U test.

## Discussion

The main finding of this single center propensity-matched retrospective cohort study was that ICU admission during medical handover did not affect resource use or clinical outcomes.

Several studies have addressed how communication errors during handover can affect clinical outcomes [8, 9, 18]. For instance, a prospective study in a tertiary ICU in Brazil showed that, among intensivists, diagnoses and goals of treatment are either not conveyed or retained in 50–60% of the cases immediately after a handover without a handover protocol, demonstrating important loss of information after transitions between staff intensivists [19]. Therefore, improve handover processes is crucial to improve patient safety, reduce medical errors and preventable adverse events [20]. Indeed, the implementation of a handover program based on a standardized oral and written handovers and communication training was associated with 23% decrease in medical-error and 30% decrease in preventable adverse events [21].

The concept that patient admission during handover time could affect urgent patient care and clinical outcomes was recently addressed in a single center retrospective cohort study with septic patients presenting to the ED during nursing handover [3]. The authors showed no significant differences in time to antibiotic administration, time to availability of serum lactate result, time to obtain blood culture and hospital mortality between patients who arrived to the ED during nursing handover time compared to those patients who did not arrive during handover time [3].

However, there are some evidence that patient care during handovers are associated with worse outcomes in surgical patients. A retrospective population-based cohort study including 313,066 patients submitted to a major surgery in Canada evaluated the effect of a complete intraoperative handover of anesthesia care from one physician anesthesiologist to another compared with no handover of anesthesia care on clinical outcomes [22]. The authors reported that patients submitted to medical transitions of care during surgery had a higher risk of adverse post-operative outcomes [22]. Another retrospective single center study found that handover of anesthetic care during cardiac surgery was associated with 43% higher risk of in hospital mortality [12]. Accordingly, patients who experience transitions of anesthesia care during surgery experience worse outcomes than those patients who did not, mostly due to information loss [22].

Our study has limitations. Our study was performed in single ICU located in a private tertiary care hospital in Brazil. Therefore, our results may not be generalizable to other ICUs localized in other developing countries, where healthcare systems and patient characteristics may vary substantially from our cohort. Secondly, we used a propensity score matching design aiming to mitigate confounding and enhance internal validity of this analysis. However, although a propensity score design helped to account for inherent differences in patient characteristics between the groups, we cannot assurance that it mitigates confounding completely [16]. Third, the handover time was arbitrarily defined as the time period between 6:30 am to 7:30 am and between 6:30 pm to 7:30 pm. Finally, it is important to highlight that our study did not investigate the impact of ICU admissions during handover on ICU staff performance, patient and staff satisfaction, patient and family experience and on the incidence of medical errors. These questions should to be further evaluated.

## Conclusion

In this propensity-matched single center retrospective cohort study, patients admitted to ICU during handover did not have higher mortality or use of resources than patients not admitted during handover. Further large-scale multicenter prospective studies are needed to improve our understanding of the association between admission during handover time and outcomes.

## Supporting information

**Fig S1.** Distribution of propensity scores along with kernel density estimates in handover and control groups before and after matching.

Treated represents handover group and control represents control group.

**Fig S2.** Absolute standardized differences comparing baseline covariates between handover group and control group patients in the unmatched and matched cohorts. White open circles represent unmatched (Before propensity score matching) and black filled circles matched (after propensity score matching) cohorts. The sixteen clinically relevant patients characteristics were entered in a logistic regression for propensity score estimation as follows: age (years), gender (0=female, 1=male), SAPS III score (points), reason for index ICU admission (0=surgical, 1=medical), index admission source (0=emergency department, 1=ward, 2=step down unit, 3=operating room/procedure, 4=others), presence of systemic hypertension (0=no, 1=yes), diabetes mellitus (0=no, 1=yes), cancer (0=no, 1=yes), congestive heart failure (0=no, 1=yes), COPD =chronic obstructive pulmonary disease (0=no, 1=yes), chronic kidney disease (0=no, 1=yes) and liver cirrhosis (0=no, 1=yes), supportive therapy during the ICU stay [need for vasopressors (0=no, 1=yes), mechanical ventilation (0=no, 1=yes), NIV=noninvasive mechanical ventilation (0=no, 1=yes) and RRT=renal replacement therapy (0=no, 1=yes)].

**Table S1.** Baseline characteristics of study participants before propensity score matching. Values represent median (IQR) or n (%). SAPS III: simplified acute physiology score III, §: scores on SAPS III range from 0 to 217, with higher scores indicating more severe illness and higher risk of death. COPD: chronic obstructive pulmonary disease, RRT: renal replacement therapy, #: another hospital and home care. * P values were calculated with the use of (a) independent t test.

## References

1. Estenssoro E, Alegria L, Murias G, Friedman G, Castro R, Nin Vaeza N, et al. Organizational Issues, Structure, and Processes of Care in 257 ICUs in Latin America: A Study From the Latin America Intensive Care Network. Crit Care Med 2017;45(8):1325–36.

2. Soares M, Bozza FA, Angus DC, Japiassu AM, Viana WN, Costa R, et al. Organizational characteristics, outcomes, and resource use in 78 Brazilian intensive care units: the ORCHESTRA study. Intensive Care Med 2015;41(12):2149–60.

3. Alsolamy S, Al-Sabhan A, Alassim N, Sadat M, Qasim EA, Tamim H, et al. Management and outcomes of patients presenting with sepsis and septic shock to the emergency department during nursing handover: a retrospective cohort study. BMC Emerg Med 2018;18(1):3.

4. Colvin MO, Eisen LA, Gong MN. Improving the Patient Handoff Process in the Intensive Care Unit: Keys to Reducing Errors and Improving Outcomes. Semin Respir Crit Care Med 2016;37(1):96–106.

5. Vincent JL, De Backer D. Circulatory shock. N Engl J Med 2013;369(18):1726–34.

6. Cavallazzi R, Marik PE, Hirani A, Pachinburavan M, Vasu TS, Leiby BE. Association between time of admission to the ICU and mortality: a systematic review and metaanalysis. Chest 2010;138(1):68–75.

7. Petersen LA, Brennan TA, O’Neil AC, Cook EF, Lee TH. Does housestaff discontinuity of care increase the risk for preventable adverse events? Ann Intern Med 1994;121(11):866–72.

8. Arora V, Johnson J, Lovinger D, Humphrey HJ, Meltzer DO. Communication failures in patient sign-out and suggestions for improvement: a critical incident analysis. Qual Saf Health Care 2005;14(6):401–7.

9. Kitch BT, Cooper JB, Zapol WM, Marder JE, Karson A, Hutter M, et al. Handoffs causing patient harm: a survey of medical and surgical house staff. Jt Comm J Qual Patient Saf 2008;34(10):563–70.

10. Ye K, Mc DTD, Knott JC, Dent A, MacBean CE. Handover in the emergency department: deficiencies and adverse effects. Emerg Med Australas 2007;19(5):433–41.

11. Kachalia A, Gandhi TK, Puopolo AL, Yoon C, Thomas EJ, Griffey R, et al. Missed and delayed diagnoses in the emergency department: a study of closed malpractice claims from 4 liability insurers. Ann Emerg Med 2007;49(2):196–205.

12. Hudson CC, McDonald B, Hudson JK, Tran D, Boodhwani M. Impact of anesthetic handover on mortality and morbidity in cardiac surgery: a cohort study. J Cardiothorac Vasc Anesth 2015;29(1):11–6.

13. Ponzoni CR, Correa TD, Filho RR, Serpa Neto A, Assuncao MSC, Pardini A, et al. Readmission to the Intensive Care Unit: Incidence, Risk Factors, Resource Use, and Outcomes. A Retrospective Cohort Study. Ann Am Thorac Soc 2017;14(8):1312–9.

14. Zampieri FG, Soares M, Borges LP, Salluh JIF, Ranzani OT. The Epimed Monitor ICU Database(R): a cloud-based national registry for adult intensive care unit patients in Brazil. Rev Bras Ter Intensiva 2017;29(4):418–26.

15. Moreno RP, Metnitz PG, Almeida E, Jordan B, Bauer P, Campos RA, et al. SAPS 3--From evaluation of the patient to evaluation of the intensive care unit. Part 2: Development of a prognostic model for hospital mortality at ICU admission. Intensive Care Med 2005;31(10):1345–55.

16. Austin PC. Statistical criteria for selecting the optimal number of untreated subjects matched to each treated subject when using many-to-one matching on the propensity score. Am J Epidemiol 2010;172(9):1092–7.

17. Austin PC. Optimal caliper widths for propensity-score matching when estimating differences in means and differences in proportions in observational studies. Pharm Stat 2011;10(2):150–61.

18. Horwitz LI, Meredith T, Schuur JD, Shah NR, Kulkarni RG, Jenq GY. Dropping the baton: a qualitative analysis of failures during the transition from emergency department to inpatient care. Ann Emerg Med 2009;53(6):701–10 e4.

19. Dutra M, Monteiro MV, Ribeiro KB, Schettino GP, Kajdacsy-Balla Amaral AC. Handovers Among Staff Intensivists: A Study of Information Loss and Clinical Accuracy to Anticipate Events. Crit Care Med 2018;46(11):1717–21.

20. Nasca TJ, Day SH, Amis ES, Jr., Force ADHT. The new recommendations on duty hours from the ACGME Task Force. N Engl J Med 2010;363(2):e3.

21. Starmer AJ, Spector ND, Srivastava R, West DC, Rosenbluth G, Allen AD, et al. Changes in medical errors after implementation of a handoff program. N Engl J Med 2014;371(19):1803–12.

22. Jones PM, Cherry RA, Allen BN, Jenkyn KMB, Shariff SZ, Flier S, et al. Association Between Handover of Anesthesia Care and Adverse Postoperative Outcomes Among Patients Undergoing Major Surgery. JAMA 2018;319(2):143–53.

